# Salty surfaces deter feeding in a blood-sucking disease vector

**DOI:** 10.1101/2021.03.22.436426

**Authors:** G Pontes, JM Latorre-Estivalis, ML Gutiérrez, A Cano, M Berón de Astrada, MG Lorenzo, RB Barrozo

## Abstract

Salts are essential nutrients required for many physiological processes, and deficient or excessive salt results in adverse health problems. Taste is the ultimate sensory modality involved in resource quality assessment, resulting in acceptance or rejection. Here, we show that detection of high-salt substrates by a salt-sensitive antennal gustatory receptor neuron, S1-GRN, results in feeding avoidance in the hematophagous bug *Rhodnius prolixus*. Knock-down of two antennal-expressed amiloride-sensitive pickpocket channel receptors (PPKs; *RproPPK014276* and *RproPPK28*) using RNA interference, prevents avoidance of bugs to high-salt substrates. Tracing antennal GRNs to the central nervous system reveals the antennal lobes as a gustatory processing center. The identification of the gustatory basis of high-salt detection in a blood feeder provides novel targets to prevent biting and feeding, as well as to promote substrate avoidance in a relevant disease vector.

**Significance Statement:** Detection of aversive gustatory stimuli induces avoidance responses in animals. Avoidance acquires particular interest if it reduces the biting rates of blood-feeding insects of medical relevance. Here we describe the molecular and physiological basis of high-salt detection in the blood-sucking disease vector *Rhodnius prolixus*. We show that detection of high-salt substrates through two PPK receptors expressed in an antennal gustatory receptor neuron produces feeding avoidance. Understanding these gustatory-driven aversive responses allows the hitherto overlooked use of gustatory molecules as a complement to known olfactory repellents.

## Introduction

Salts are vital for physiological processes. Sodium and potassium chloride regulated at adequate levels are crucial for maintaining electrolyte homeostasis and neuronal transmission. Deficient or excessive salt results in adverse health effects. Consequently, in most animals studied so far, the assessment of salt concentration drives acceptance or avoidance behavioral responses (1, 2). For example, low-salt (< 0.1 M) triggers insect feeding and oviposition, whereas high-salt (> 0.1 M) promotes avoidance and prevents egg-laying (3–9). Salts are detected by the taste system, a sensory modality that helps animals make predictions about the value of a potential food source, *i.e.* whether beneficial or harmful. Found on the surface of the body of insects (1), taste sensilla are the structural units of this system that house gustatory receptor neurons (GRNs). GRNs carry specific receptors on their dendritic membrane that favor the detection of different types of gustatory stimuli (10). Upon GRN stimulation, a message is sent to the brain through action potentials and, consequently, the appropriate behavior is set into motion: ovipositing or not, biting or not, eating or not.

Despite the relevance of salts to life, the molecular and physiological mechanisms underlying salt perception and processing by animals are barely understood (11, 12). Degenerin-epithelium sodium channels (DEG/ENaC) have been related to salt sensing in mammals (13). Insect pickpocket channel receptors (PPK) belong to the DEG/ENaC family and were reported to detect salt in *Drosophila melanogaster*, among other functions (14). They constitute a large family of ligand-gated membrane proteins, voltage-insensitive ion channels and some of them are highly sensitive to amiloride (AMI) blockade (15, 16). The *DmelPPK11* and *DmelPPK19* genes, for example, were related to low-salt detection in the fruit fly *D. melanogaster* (4). Besides, fruit fly *DmelPPK28* and its mosquito orthologue *AaegPPK301* (*Aedes aegypti)* were identified as osmolarity sensors because the gustatory receptor neurons (GRNs) expressing them respond to hypo-osmotic solutions (5, 17, 18). High-salt sensing in *D. melanogaster*, on the other hand, occurs through specific GRNs that also detect bitter compounds (19) and a population of glutamatergic neurons expressing *DmelPPK23* (11). However, the detection of salt is complex and, in addition to PPKs, other receptors also participate, such as such *IR76b, IR94e, sano* (11, 20, 21).

In blood-feeding insects, salts such as NaCl and KCl were demonstrated to be relevant gustatory cues in a feeding context (22, 23), probably because salts are essential components of blood of vertebrates. Besides, salts are found on human skin as part of sweat, product of the eccrine glands excretion (24). Salt concentration modulates the behavior of the hematophagous bug *Rhodnius prolixus*, triggering preference or avoidance responses. For example, they avoid surfaces coated with high-NaCl or -KCl (1 M) (25, 26), preferring instead surfaces impregnated with salt concentrations similar to those of the human sweat (0.1 M) (24, 25). Furthermore, *R. prolixus* accepts or rejects a feeding solution depending on its NaCl or KCl concentration (3, 8). Many molecules produce insect avoidance due to the negative impact they may have on their physiology, *i.e.* toxic or even lethal effects (27–31). Therefore, biting and feeding decisions depend strongly on the assessment of the chemical qualities of the food source (29).

Triatomine insects (Hemiptera: Reduviidae), commonly known as kissing bugs, are vectors of the protozoan parasite *Trypanosoma cruzi,* the causative agent of Chagas disease, to humans. Whereas Chagas disease is considered endemic to Mexico, Central and South America, it has become a global health problem affecting an increasing number of non-endemic countries (32, 33). In the United States of America, the Center for Disease Control and Prevention (CDC) considered the Chagas disease as one of the five parasitic neglected infections to be targeted as priority for public health (34).

We investigated the basis of high-salt perception in the kissing bug *R. prolixus* by combining behavioral, pharmacological, electrophysiological, neuroanatomical and molecular approaches. Our work shows that kissing bugs detect NaCl and KCl with the antennae, and that high-salt doses on the bite site prevent feeding. Salt-sensitive gustatory receptor neuron 1 (S1-GRN) requires two PPKs, *RproPPK014276* and *RproPPK28*, for salt detection. Besides, the activation of S1-GRN is responsible for triggering high-salt avoidance. Moreover, we show that antennal GRNs project to the antennal lobes (a brain region mostly dedicated to the processing of olfactory information in insects), where the antennal gustatory information seems to be processed in *R. prolixus*.

## Results

### Salty bite sites prevent feeding

Kissing bugs assess the quality of a potential food source by tasting the host skin (31). Consequently we asked whether detection of high-salt over the bite site would affect feeding on an otherwise appetitive solution (AS; 0.001 M ATP in 0.15 M NaCl). To do this, the membrane of the artificial feeder mimicking the host skin was coated with different concentrations of NaCl (Na) or KCl (K): no-Na/no-K, low-Na/low-K, mid-Na/mid-K or high-Na/high-K (Figure 1A). Note that no-Na and no-K represent control groups, in which the membrane was impregnated with distilled water instead of a salt solution.

**Figure 1.**
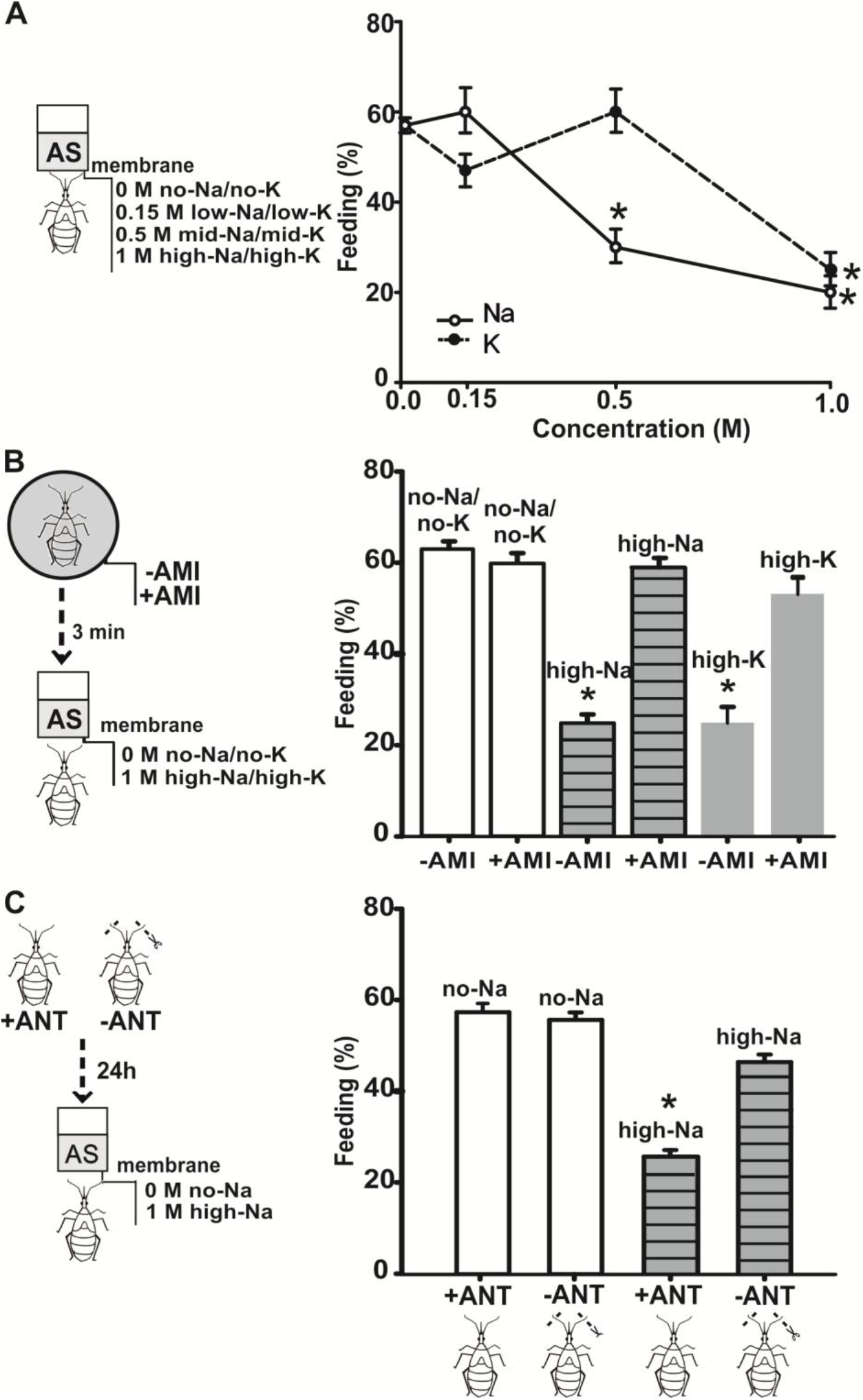
Figure 1- High-salt bite sites trigger feeding avoidance Insect feeding response to an appetitive solution (AS) when the bite membrane of the artificial feeder is coated with NaCl (Na) or KCl (K). **(A)-** Salt concentration in the bite membrane affects feeding. The bite membrane was coated with different concentrations of Na/K: no-Na/no-K (0 M), low-Na/low-K (0.15 M), mid-Na/mid-K (0.5 M) or high-Na/high-K (1 M). Dots represent the percentage of insects that fed. Insects avoided feeding after contacting membranes coated with mid-Na or high-Na/high-K. Asterisks indicate, for each salt, significant differences across concentrations (Pearson *X*^2^, *k* = 3, α’ = 0.016, p < 0.01). Twenty and thirty replicates were carried out for Na and K concentrations, respectively. **(B)-** Amiloride blocks high-salt avoidance. Insects were treated for 1 minute with 0.001 M amiloride (+AMI) or distilled water (-AMI). Three minutes later insects were exposed to membranes coated with no-Na/no-K (0 M) or with high-Na/high-K (1 M). Each bar represents the percentage of insects that fed. Amiloride blocked high-Na/high-K perception and insects fed over high-Na/high-K membranes. Asterisks indicate, for each salt, significant differences between +AMI *vs*. -AMI (Pearson *X*^2^, *k* = 3, α’ = 0.016, p < 0.01).Thirty replicates were carried out for each experimental condition. **(C)-** High-salt avoidance begins at the antennae. The tip of distal antennal flagellomeres were ablated (-ANT) or kept intact (+ANT). Twenty four hours later insects were exposed to membranes coated with no-Na (0 M) or with high-Na (1 M). Each bar represents the percentage of insects that fed. In the absence of the taste sensilla of the distal antennal flagellomeres, insects fed over high-Na coated membranes. Asterisk indicates significant differences between-ANT *vs*. +ANT (Pearson *X*^2^, *k* = 3, α’ = 0.016, p < 0.01).Thirty replicates were carried out for each treatment.

About 60 % of the insects fed over membranes coated with no-Na/no-K or low-Na/low-K (Figure 1A). No significant differences between responses to no-Na/no-K and low-Na/low-K were detected (Na: χ^2^ = 0.05, p = 0.81; K: χ^2^ = 0.61, p = 0.43). In contrast, a significant reduction in the percentage of insects feeding on the AS was observed when the bite membrane was coated with mid-Na (no-Na, low-Na *vs.* mid-Na, χ^2^ =5.98, p = 0.01; α’ = 0.016) or high-Na/high-K (no-Na, low-Na *vs.* high-Na, χ^2^ = 8.28, p = 0.004; no-K, low-K *vs.* high-K, χ^2^ = 6.04, p = 0.01; α’ = 0.016).

These results show that perception of high-Na or high-K on the bite site produces feeding avoidance in *R. prolixus*.

### High-salt avoidance is impaired by amiloride blockade

DEG/ENaCs, including insect PPKs receptors can be blocked by amiloride (4, 18, 35, 36). Therefore, we examined whether this drug could block the detection of high-salt during bite site assessment by kissing bugs. To test this, insects were exposed to amiloride (+AMI) or distilled water (-AMI). Following the treatment, feeding responses to the AS on coated membranes with no-Na/no-K or a high-Na/high-K were evaluated (Figure 1B).

The −AMI insects consistently avoided feeding if exposed to high-Na/high-K membranes (−AMI no-Na/no-K *vs.* −AMI high-Na, χ^2^ =5.88, p = 0.01; −AMI no-Na/no-K *vs.* −AMI high-K, χ^2^ =6.45, p =0.01; α’ = 0.016). However, +AMI treated insects fed over high-Na/high-K coated membranes (+AMI no-Na/no-K *vs.* +AMI high-Na χ^2^ =0.01, p = 0.9; +AMI no-Na/no-K *vs.* +AMI high-K χ^2^ =0.24, p =0.62; α’ = 0.016). As expected, insects fed normally on no-Na/no-K membranes regardless of the treatment applied (-AMI no-Na/no-K *vs. +*AMI no-Na/no-K χ^2^ = 0.05, p = 0.81; α’ = 0.016). These results show that amiloride blocks the perception of high-salt present over a feeding surface in *R. prolixus*.

### High-salt taste detection begins in the antenna

We asked, next, whether high-Na/high-K avoidance was triggered by antennal gustatory neurons. We hypothesized that ablation of the taste sensilla present on the tip of the distal flagellomere would impede high-salt sensing on surfaces, as occurs for bitter substances (31). Consequently, we expected that ablated insects would feed over a high-salt coated membrane. To test this, the tip of the distal flagellomere of both antennae was either cut (-ANT) or kept intact (+ANT). Then, we tested the feeding response to the AS of −ANT and +ANT groups, exposed to either a no-Na or high-Na coated membranes (Figure 1C).

As already shown, the presence of high-Na over the membrane prevented feeding on intact animals (+ANT no-Na *vs.* +ANT high-Na, χ^2^ = 5.55, p = 0.01; α’ = 0.016). Conversely, this avoidance vanished when the distal flagellomeres were ablated (-ANT no-Na *vs.* −ANT high-Na, χ^2^ = 0.6, p = 0.43; α’ = 0.016). Note that cutting off the tip of the distal flagellomeres imposed no negative effects on insect feeding performance (-ANT no-Na *vs.* +ANT no-Na, χ^2^ = 0.60, p = 0.43; α’ = 0.016).

These results indicate that the detection of high-Na on surfaces occurs in taste sensilla located on the distal antennal flagellomeres of *R. prolixus*.

### Two antennal GRNs detect salts

Then, we examined the presence of salt-sensitive GRNs in the 4 taste sensilla of the tip of the distal antennal flagellomeres (Figure 2A). Single-sensillum taste recordings on these sensilla (shown in Figure 2A) revealed 2 GRNs sensitive to Na and K, S1-GRN and S2-GRN. Both GRNs were easily distinguished by the amplitude and waveform of their spikes (Figure 2A). S1-GRN and S2-GRN increased their firing rate with salt concentration (Figure 2B, 2C). Whereas S1-GRN responded to both salts, S2-GRN was only responsive to Na (Figure 2C). The temporal firing profiles of both neurons also showed notable differences. S1-GRN responded to Na/K in a phasic-tonic manner (Figure 2D), while S2-GRN fired tonically upon Na stimulation (Figure 2E).

These results reveal that the 4 taste sensilla of the tip of the distal flagellomeres house at least two GRNs tuned either to Na and K or Na alone, and that the salt receptors or transduction mechanisms of each GRN are different.

**Figure 2.**
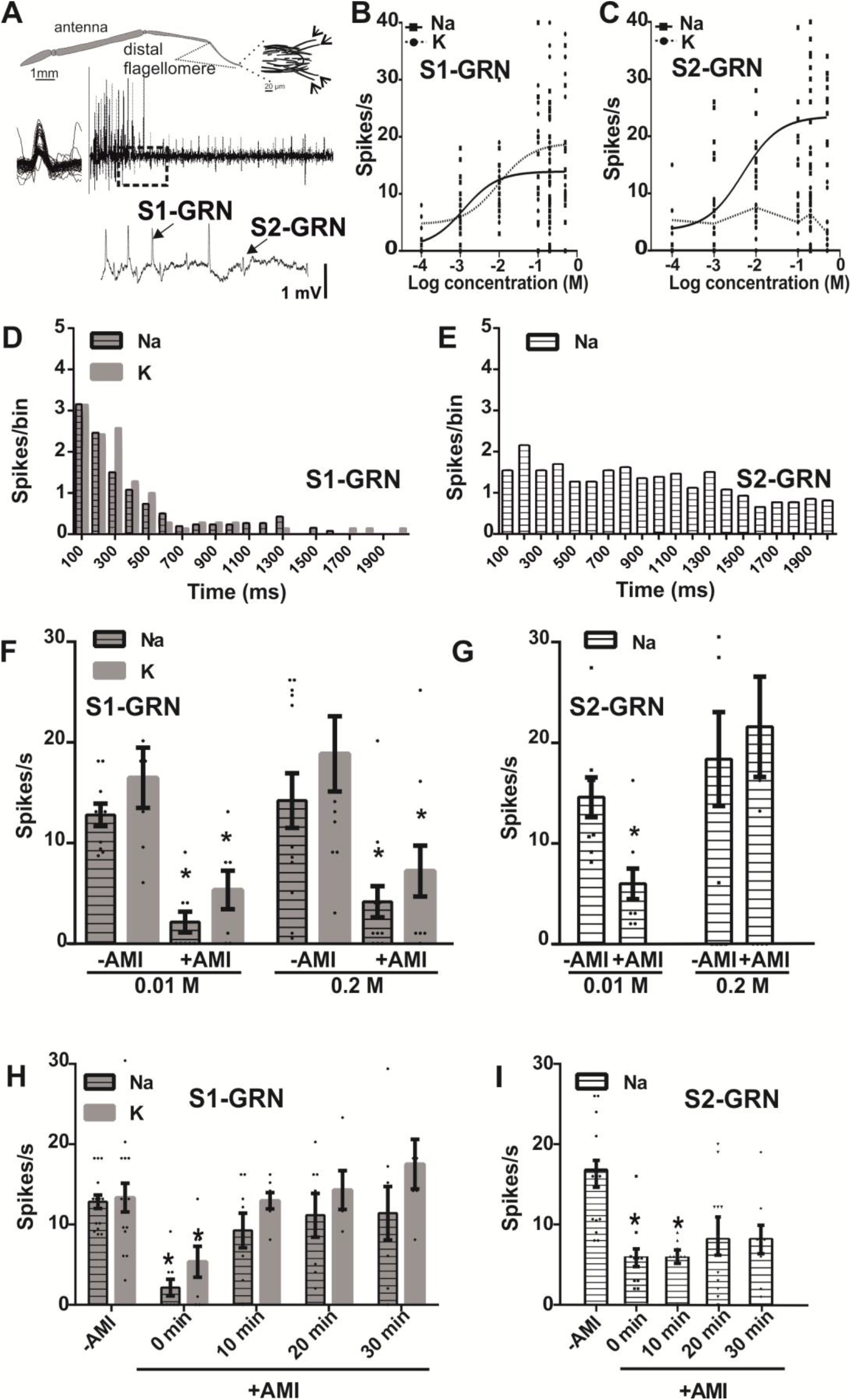
Antennal gustatory receptor neurons involved in salt detection. **(A)**- Scheme of the antenna of *R. prolixus* and of the end of the distal flagellomere. The arrows indicate the 4 recorded taste sensilla. Example of spike discharges of the GRNs housed in a taste sensillum. A typical firing response to 0.01 M Na is shown. Two salt-responsive GRNs were identified, S1-GRN and S2-GRN. As shown in the magnified area of the single-sensillum recording, S1-GRN showed action potentials of higher amplitude than S2-GRN. **(B-C)**- Antennal GRNs respond to salts. Scatter plots and fitted curves of spike frequencies are shown. S1-GRN and S2-GRN responded to Na in a dose-dependent manner, but only S1-GRN responded also to K. Between six and twenty-seven sensilla were tested for each salt concentration. **(D-E)-** Differing dynamics of the temporal pattern of GRNs responses. Peri-stimulus histogram of S1-GRN and S2-GRN in response to 0.01 M Na or K. Each bar represents the mean number of action potentials recorded for each 100 ms bin. S1-GRN showed a phasic-tonic response profile, which was similar for both salts. In contrast, S2-GRN showed a tonic response pattern. Seven to thirteen taste sensilla were tested for each GRN type. **(F-G)-** Amiloride blocks GRNs responses to salts. Taste sensilla were treated with 0.001 M AMI (+AMI) or with distilled water (-AMI). Scatter plots are shown and bars represent the mean of spike frequency (mean ± s.e.m) of GRNs in response to 0.01 and 0.2 M Na or K. The firing response of S1-GRN was significantly reduced by +AMI to both salts and at the two concentrations. In contrast, the firing activity of S2-GRN was only blocked by +AMI at 0.01 M Na. Asterisks indicate statistical differences between +AMI and -AMI (Wilcoxon test, α = 0.05, p < 0.03) for each salt. Nine to thirteen taste sensilla were tested per condition. **(H-I)-** Differing recovery time of GRNs responses after amiloride blockade. Taste sensilla were applied with AMI as before (F-G). After +AMI application (0 min), and subsequently at 10 minute intervals, the firing rate of GRNs was recorded. -AMI, represents a control of the response without AMI application. Scatter plots are shown and bars represent the mean of spike frequency (mean ± s.e.m) of GRNs in response to 0.01 M Na or K. The recovery time of both GRNs is different. S1-GRN firing response recovered faster than S2-GRN. Asterisks indicate, for each salt, statistical significant differences between the -AMI group and the +AMI groups at each post-application time (Wilcoxon test, *k* = 4, α’ = 0.0125, p < 0.002). Six to ten taste sensilla were tested for each GRN type.

### Amiloride interferes with salt detection

Figure 1B showed that AMI blocks the perception of high-Na/high-K. Therefore, we expected that +AMI treatment would have an effect on the firing activity of GRNs. To test this, taste sensilla were treated with +AMI, and subsequently, the response of both S1-GRN and S2-GRN to Na or K was recorded (Figure 2F-2I).

AMI blocked the activity of S1-GRN and S2-GRN by significantly decreasing their firing rate to salts (Figure 2F, 2G). However, while the +AMI blocking effect on S1-GRN responses was independent of the Na/K concentration (Figure 2F, 0.01 M Na: W = −45 p =0.002; 0.2 M Na: W = −83, p = 0.0009; 0.01 M K: W = −28, p =0.008; 0.2 M K: W = −37, p = 0.03), it only reduced S2-GRN responses at the lowest Na concentration tested (Figure 2G, 0.01 M Na: W = −68, p = 0.002; 0.2 M: W = 14, p = 0.28).

The recovery time of GRNs following +AMI application was also analyzed based on the response profiles (Figure 2H, 2I). The firing activity of S1-GRN to Na/K was significantly reduced immediately after AMI application (Figure 2H, −AMI vs. +AMI 0 min: Na, W = −45, p = 0.002; K: W = −28, p = 0.007; α’ = 0.0125) and recovered 10 minutes later (-AMI vs. +AMI 10 min: Na, W = −18, p = 0.07; K: W = −10, p = 0.21; −AMI vs. +AMI 20 min: Na, W = −8, p = 0.29; K: W = −3, p = 0.41; −AMI vs. +AMI 30 min: Na, W = −10, p = 0.26; K: W = 4, p = 0.41; α’ = 0.0125). A decreased response to Na following AMI exposure was also evident for S2-GRN which recovered after a similar interval (Figure 2I, −AMI *vs*. 0 min or 10 min, W = −68, p= 0.002, W = −42, p = 0.006; α’ = 0.0125). Even though the firing response of S2-GRN seemed lower 20 minutes after AMI exposure, the decrease was not statistically significant (-AMI vs. +AMI 20 min, W = −37, p = 0.03; −AMI vs. +AMI 30 min, W = −29, p = 0.04; α’ = 0.0125).

These data show consistently that amiloride blocks salt detection in the GRNs of the antennae. They also show that the effect of AMI on both GRNs is markedly different.

### Antennal GRNs project into the ipsilateral antennal lobe

Next, we traced the neurons housed in the 4 taste sensilla of the distal flagellomeres (Figure 2A), in order to examine their projections into the central nervous system. Initially, backfills with neurobiotin were attempted in a single cut sensillum. Two out of 19 preparations were successful, allowing us to trace their projections to ipsilateral the antennal lobe (AL) (Figure 3A, 3B). In order to reinforce this initial evidence, we subsequently carried out additional mass fills by distally cutting all 4 sensilla of a single antenna simultaneously, and exposing them to rhodamine. In order to assure that only neurons of these taste sensilla were stained, the 4 sensilla of one antenna were cut, while the same sensilla of the other antenna were kept intact. Thereafter, the sensilla of both antennae, *i.e.* cut and intact sensilla, were placed in glass capillaries containing rhodamine. Seven out of 12 preparations of cut tip sensilla were successfully stained, confirming that neurons housed in these sensilla send projections to the ipsilateral AL.

**Figure 3.**
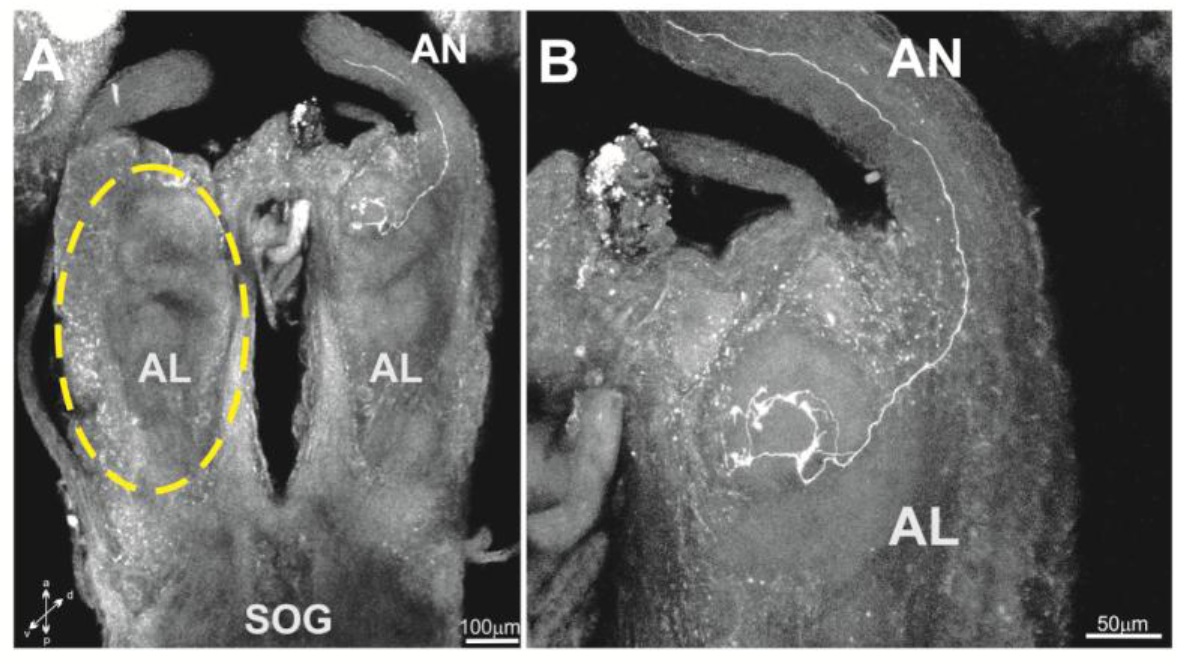
Antennal GRNs project into the antennal lobes. **(A)-** Example of a backfill of a single taste sensillum with neurobiotin. A single GRN became stained and was found to project into the ipsilateral AL. The dashed line delimits the contralateral AL. **(B)-** Enlargement of the AL showing the arborizations of the GRN stained in (A). Antennal lobe: AL, SOG: subesophageal ganglion, AN: antennal nerve.

Stained reparations based on both techniques allowed determining that the axons of antennal GRNs arborise mainly at the anterior and ventral side of the ipsilateral AL (Figure 3A, 3B). This neuropil was the only brain structure stained in all 9 successful preparations.

These results suggest that ALs are the primary center for processing antennal gustatory inputs.

### Two PPK salt receptor candidates

Next, salt receptor candidates were searched for in the genome of *R. prolixus*. We focused on PPKs because of their known role in salt sensing in insects (4, 5, 11). The genome of *R. prolixus* revealed 10 PPK protein sequences with a length between 337 and 564 amino acids (37). These PPK protein sequences presented the amiloride-sensitive sodium domain (Latorre-Estivalis *et al.*, submitted). Among them, two PPKs were chosen as salt receptor candidates, the *RproPPK014276* (identified in VectorBase as RPRC014276) and the *RproPPK28* (identified in VectorBase as RPRC000471). The choice of these candidates was based on their clustering with *D. melanogaster* or *Ae. aegypti* PPK receptors (*DmelPPK19, DmelPPK28* and *AaegPPK301*) with a role in salt sensing or in detection of hipo-osmotic solutions (4, 5, 17, 18). The *RproPPK014276* clustered with the *DmelPPK19* and belong to the PPK subfamily III, and the *RproPPK28* which is orthologous of *DmelPPK28* and belong to the PPK subfamily V (Latorre-Estivalis *et al*. submitted). Besides, *RproPPK28* also clustered in the same clade with *Ae. aegypti AaegPPK301*. Therefore, we checked for the expression of *RproPPK014276* and *RproPPK28* in the distal antennal flagellomeres of *R. prolixus*. The RT-qPCR products found after 32-34 (*RproPPK014276)* and 29-32 threshold cycles (*RproPPK28)*, confirmed their presence among the antennal transcripts (Figure 4A).

**Figure 4.**
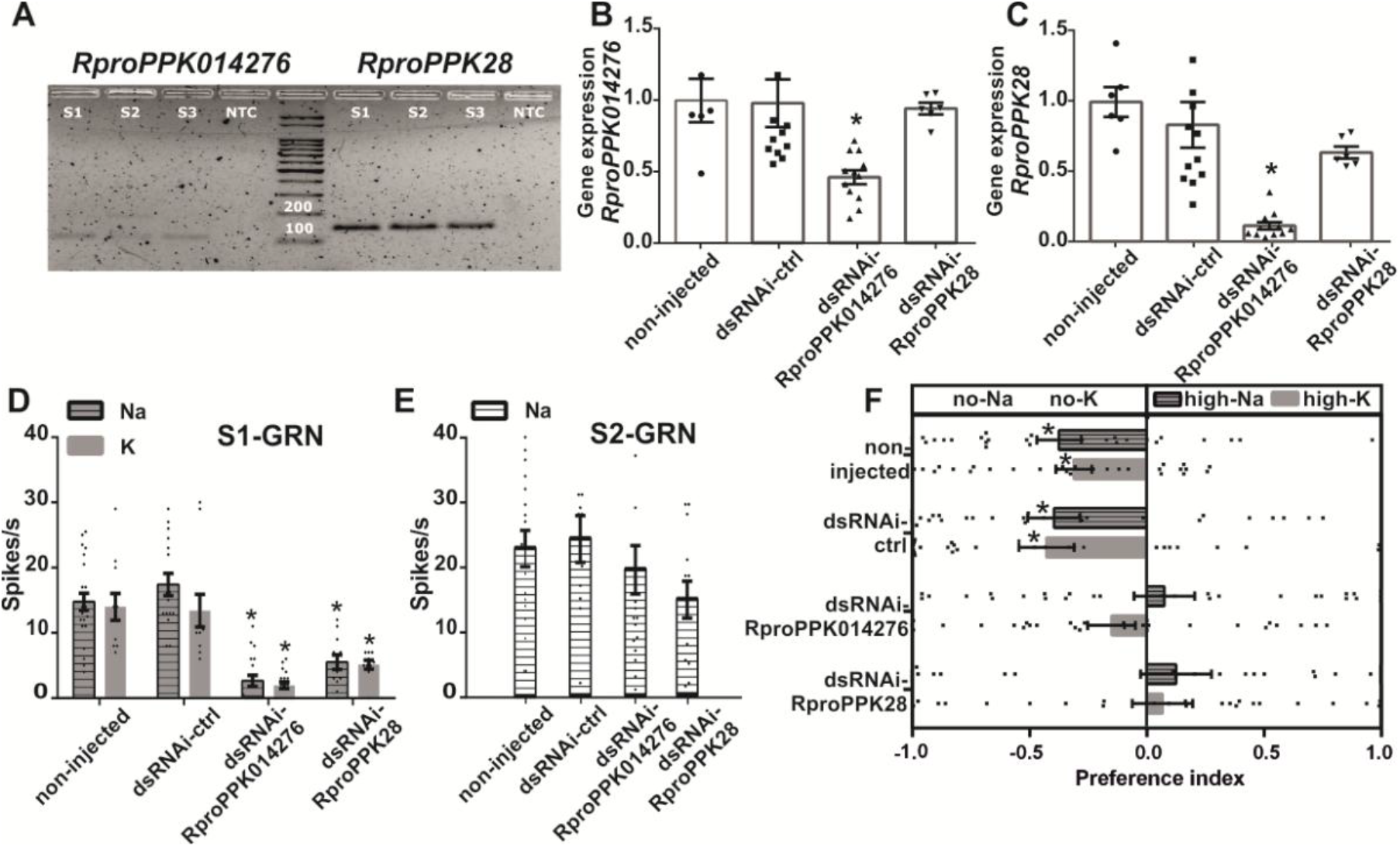
High-salt avoidance depends on *RproPPK014276* and *RproPPK28*, the functional salt receptors of S1-GRN. **(A)-***RproPPK014276* and *RproPPK28* express in the distal antennal flagellomeres. Electrophoresis of qPCR products of 3 replicates (S1, S2, S3) and a non-template control (NTC). *RproPPK014276* and *RproPPK28* qPCR products were detected after threshold cycles of 32-34 and 29-32, respectively, confirming their expression in the antenna. **(B-C)-** Transcript levels of *RproPPK014276* and *RproPPK28* decay following RNA interference. Scatter plots are shown and bars represent the relative transcript levels (mean ± s.e.m) of *RproPPK014276* (B) and *RproPPK28* (C) in non-injected and dsRNAi-injected groups. The expression levels of *RproPPK014276* and *RproPPK28* decayed significantly only when the corresponding dsRNAi was injected. Selective knock-down of the gene of interest was also confirmed, as the dsRNAi-*RproPPK28* group showed similar transcript levels for the *RproPPK014276* gene as those observed in control groups (B, last bar). Similarly, the dsRNAi-*RproPPK014276* group showed similar transcript levels for *RproPPK28* as did the control groups (C, last bar). Asterisks denote significant differences across groups (Kruskal-Wallis test, Dunn’s *post hoc* comparisons, α = 0.05, p < 0.001). Six to twelve replicates were carried out per treatment. **(D-E)-***RproPPK*s knock-down negatively affects salt detection. Spike frequencies of S1-GRN and S2-GRN to 0.01 M Na or K in non-injected and dsRNAi-injected groups. Scatter plots are shown and bars represent the mean spike frequency recorded for each condition (mean ± s.e.m). Both, *RproPPK014276 and RproPPK28* are required for proper S1-GRN response to Na or K (D). In contrast, *RproPPK*s silencing had no effect on the responses of S2-GRN to Na (E). Asterisks denote, for each salt, statistical differences between non-injected and dsRNAi-injected groups (Kruskal-Wallis test, Dunn’s *post hoc* comparisons, α = 0.05, p < 0.0001). Twelve to twenty-three sensilla for each experimental condition were tested. **(F)-** Loss of high-salt avoidance after *RproPPK*s silencing. Preference of kissing bugs was tested in a simultaneous two-choice arena, whose half zone was coated with high-Na/high-K (1 M) and the other half with no-Na/no-K (0 M, distilled water). Scatter plots are shown and bars represent the mean preference index (PI) (mean ± s.e.m), 0 denoting equal time spent at each zone of the arena, −1 denoting avoidance for the high-Na/high-K zone, and 1 denoting preference for the high-Na/high-K zone. Non-injected and dsRNAi-ctrl groups avoided high-Na/high-K coated substrates. dsRNAi-*RproPPK014276* and dsRNAi-*RproPPK28* groups showed a random behavior with no preference for any zone of the arena. Asterisks denote, for each salt, statistical differences between the PI and 0 for each experimental group (*t*-test, α = 0.05, p < 0.001). Thirty replicates were carried out for each experimental condition. Non-injected: intact control group, dsRNAi-ctrl: ds-RNA against ß-lact, dsRNAi-*RproPPK014276*: ds-RNA against *RproPPK014276*, dsRNAi-*RproPPK28*: ds-RNA against *RproPPK28*.

Based on these findings, *RproPPK014276* and *RproPPK28* were candidates as salt receptors in *R. prolixus*.

### *RproPPK014276* and *RproPPK28* genes can be knocked down

Next, RNA interference experiments were carried out in order to study the role of *RproPPK014276* and *RproPPK28* in high-salt detection. Four experimental groups of insects were prepared: non-injected, dsRNAi- ctrl, dsRNAi-*RproPPK014276* and dsRNAi-*RproPPK28*. dsRNAi- ctrl consisted of a control group of ds-RNA injection (see Materials and Methods). Effective knock-down of transcript levels of *RproPPK014276* and *RproPPK28* was first assessed by RT-qPCR (Figure 4B, C).

A significant reduction of transcript levels for *RproPPK014276* (Figure 4B, H = 19.6, p = 0.0002) and *RproPPK28* (Figure 4B, H = 25.2, p = 0.0001) was well established with respect to non-injected and dsRNAi-ctrl groups (*post hoc* Dunn’s comparisons, p < 0.05). Furthermore, we checked that gene knock-down was specifically targeted to the mRNA of interest in each ds-RNAi group. For this, we checked that the expression of *RproPPK28* was not affected when knocking down *RproPPK014276* and *vice versa*, as shown in Figure 4B, 4C (see last bars).

### Knock-down of *RproPPK014276* or *RproPPK28* abolishes salt detection only in S1-GRN

Then, we examined whether knock-down of *RproPPK014276* and *RproPPK28* altered the response of S1-GRN and S2-GRN to Na or K. Consequently, the firing activity of S1-GRN and S2-GRN was examined on the 4 taste sensilla (Figure 2A) of non-injected and dsRNAi-injected insects (Figure 4D, 4E).

The spike frequency of S1-GRN to Na or K was significantly decreased (Figure 4D, Na: H = 44.2, p < 0.0001; K: H = 38.2, p < 0.0001) in dsRNAi-*RproPPK014276* and in dsRNAi-*RproPPK28* groups with respect to non-injected and dsRNAi-ctrl ones (*post hoc* Dunn’s comparisons, p < 0.05). In contrast, the firing rate of S2-GRN to Na showed no significant difference across groups (Figure 4E, H = 5.2, p = 0.15).

These results show that *RproPPK014276* and *RproPPK28* participate in Na and K detection in S1-GRN. Furthermore, the results reveal that a decreased expression of either of these genes suppresses the response to Na or K, demonstrating that both PPKs are necessary for S1-GRN to detect salts.

### *RproPPK014276* and *RproPPK28* are responsible for high-salt avoidance

Next, we examined whether *RproPPK014276* and *RproPPK28* were involved in the avoidance of salty substrates. Kissing bugs avoid high-Na/high-K substrates if an alternative clean substrate choice is available (25). Therefore, we tested the preference of knocked down and control insects for substrates coated with high-Na/high-K *vs.* no-Na/no-K. To test this, half of substrate of a two-choice arena was coated with high-Na/high-K while the other half was kept salt free (Figure 4F). As expected, non-injected and dsRNAi-ctrl insects avoided the high-Na/high-K zone of the two-choice arena (non-injected, Na: t = −3.93, p = 0.0005, K: t =−4.01, p = 0.0004; dsRNAi-ctrl, Na: t = −3.59, p = 0.0013, K: t = −3.60, p =0.0013). In contrast, dsRNAi-*RproPPK014276* and dsRNAi-*RproPPK28* injected insects showed no significant preference for a zone of the arena, *i.e.*, they lost their avoidance of the high-Na/high-K zone (dsRNAi-*RproPPK014276*, Na: t =0.60, p = 0.55, K: t =−1.45, p = 0.15; dsRNAi-*RproPPK28*, Na: t=0.57, p = 0.57, K: t =0.53, p =0.60).

These results show that misexpression of *RproPPK014276* or *RproPPK28* prevents high-Na/high-K sensing, and therefore high-salt avoidance. Furthermore, the expression of both genes is necessary to signal the presence of high-salt and, ultimately, to trigger the avoidance behavior to salty substrates.

## Discussion

Innate avoidance is triggered when a threat is detected through interaction with deterrent sensory inputs. Our study reveals the molecular and physiological basis of high-salt taste detection in a blood-sucking insect of great epidemiological interest. We have shown evidence that allowed us to characterize the response of *R. prolixus* to high-salt, and to identify the sensory organs, chemosensory neurons, molecular taste receptors mediating high-salt detection. Besides we uncovered the downstream brain regions for antennal taste processing.

### High-salt taste detection on the bite site prevents feeding

*R. prolixus* does a gustatory evaluation of the feeding and walking substrate (25, 26, 31). Here, we showed that high doses (1M) of both NaCl and KCl negatively interfered with feeding. In contrast, low-salt doses (0.15M) did not modulate the feeding behavior of kissing bugs, not enhancing biting/feeding probability relative to a salt-free bite site (no-Na/no-K). Detection of high-salt content on the bite site begins at the antenna level (Figure 1B). In absence of the distal antennal flagellomeres, feeding avoidance of kissing bugs over high-salt substrates vanished. Likewise, feeding inhibition triggered by detection of bitter compounds in *R. prolixus* was reverted when the distal ends of both antennae were absent (31). Nevertheless it cannot be ruled out that other taste sensilla present, for example on the tarsi, can also detect gustatory stimuli (38). However, it seems that the antennae are the key taste organs to signal the presence of negative inputs during bite site evaluation in *R. prolixus*. High-salt content elicits food avoidance on most species studied so far (4, 7, 11, 39). Yet, food avoidance triggered by high-salt detection on the bite site is shown here for the first time in a hematophagous insect.

### Antennal GRNs involved in salt detection

Two GRNs housed in the taste sensilla of the distal flagellomeres of *R. prolixus* are responsible for antennal salt detection. Although both were activated by NaCl, only S1-GRN1 was also sensitive to KCl. S1-GRN1 and S2-GRN2 showed clear physiological differences in the temporal firing patterns in response to salts. Whereas S1-GRN showed phasic-tonic responses, S2-GRN2 exhibited tonic responses when stimulated with NaCl. The phasic component of the response of receptor neurons may provide significant information about stimulus intensity and quality (40–42). Besides, a defined peak of impulse frequency enables the animal to recognize the onset of stimulation (43). The existence and relevance of a spatiotemporal code across GRNs, likely signaling the quality and intensity of the taste stimulus has been shown by several studies (44–47). Thus, S1-GRN could quickly warn the insect about the presence of a noxious salty stimulus, whereas a tonic response such as in S2-GRN could indicate its persistence.

Both GRNs were sensitive to amiloride blockade, although effects were different for each neuron type. These findings demonstrated the existence of at least two different amiloride-sensitive salt receptors in each GRN. Looking at the response of S1-GRN to the highest dose of salt tested allows inferring a kind of non-competitive blockade by amiloride, as amiloride still blocked the responses by S1-GRN but not by S2-GRN. Amiloride blockade at the receptor level translated into absence of behavioral responses to high-salt content, and consequently, *R. prolixus* feed over a high-salt bite site. Similarly, avoidance of high-salt substrates was impaired when kissing bugs were treated with amiloride and tested in a dual-choice preference arena (25). Susceptibility of salt receptors to amiloride blockade has also been described in *D. melanogaster* (4) and rodents (13, 48–50). Yet and contrary to what we found in this work, only salt receptors narrowly tuned to NaCl were inhibited by amiloride in rodents, but not those responsive to both salts, NaCl and KCl (13, 48–50).

### Two PPKs are required for high-salt taste

Ten PPKs genes had been previously identified to be expressed in the antennae of *R. prolixus* (37). However, the functional role of PPKs in hematophagous insects had not been studied until recently (5, and this work). As a first approach, we focused on *RproPPK014276* and the *RproPPK28* because they clustered with *DmelPPK19* (from *D. melanogaster*), and *DmelPPK28* and *AaegPPK301* (from *D. melanogaster* and *Ae. aegypti*, respectively), respectively. Notably *DmelPPK19* has been related to ion detection (4). Following disruption of *DmelPPK19* flies failed to recognize low NaCl or KCl concentrations (4). *DmelPPK28* was associated instead to detection of hypo-osmotic solutions in flies (17, 18). Recently, *AaegPPK301* was found to be sensitive to low-salt and required for the initiation of egg-laying by females of the mosquito *Ae. aegypti* (5).

*DmelPPK19* has also been related to high-salt detection in *D. melanogaster* (4). As predicted for known DEG/ENaC and PPKs sequences, it was shown that *R. prolixus* PPKs present two transmembrane domains, a cysteine-rich domain and a Deg motif, and revealed the presence of an amiloride-binding domain next to the pore region (Latorre-Estivalis *et al*., submitted). But even more important, we showed that following *RproPPK014276* and *RproPPK28* knock-down, avoidance for high-Na/high-K substrates in *R. prolixus* vanished. Notably, to generate high-salt avoidance both PPKs are necessary, as inactivating a single PPK was enough to disrupt the behavioral responsiveness to high salt. Indeed Ben-Shahar *et al*. (15) proposed that PPKs form homo- or heteromeric protein complexes of three subunits (or multiples of three). Our results suggest that PPKs could form heteromeric protein complexes in kissing bugs gustatory receptor neurons.

### One GRN and two PPKs are responsible for high-salt avoidance

Knocking down the expression of either *RproPPK014276* or *RproPPK28*, induced an abrupt decay of S1-GRN firing responses to NaCl and KCl. As a consequence, insects showed a random behavior on the two-choice preference arena, loosing avoidance for a high-salt substrate. Interestingly, S2-GRN does also respond to NaCl but their activity was not affected by PPKs silencing, indicating that S2-GRNs express a different type/s of salt receptor/s than S1-GRNs. As mentioned above, receptors other than PPKs seem to be involved in salt sensing in *D. melanogaster,* as for example *IR76b*, *IR94e* and *sano* (9, 11, 20, 21). More than one neuron/cell seems to be sensitive to salts in all animals studied so far (4, 11, 13, 39, 49, 51). Interestingly, Masagué *et al*. (25) demonstrated previously, through discriminative experiments, that *R. prolixus* cannot distinguish between NaCl and KCl. After this work, we can infer that NaCl or KCl cannot be discriminated because their detection depends on two PPKs expressed by the same neuron. Therefore messages conveying information about the detection of aversive salt concentration on surfaces -without discrimination across salts-would be driven to the brain through S1-GRN. However, kissing bugs can distinguish between salty and bitter tastes (25). Whether bitter tastes are also detected by the same GRN that detects high-salt or not remains to be elucidated in *R. prolixus* (11, 19, 51). Two salt-sensitive GRNs (*i.e.* GR66a-GRNs and PPK23-GRNs) have been associated with high-salt detection by *D. melanogaster* (11). These high-salt sensing GRNs of fruit flies showed to be ion non-selective, in contrast to the low-salt sensing GRN (*i.e.* GR64f-GRN) which was specifically tuned to NaCl (11). Here, we have shown that S1-GRN activation, depending on *RproPPK014276* and *RproPPK28* expression, provides kissing bugs with useful information about the deterrent saltiness of a substrate.

### Antennal gustatory inputs are processed in the ALs

Antennal receptor neurons send their axons *via* the antennal nerves to the deutocerebrum, which is composed by two parts: the antennal lobe (AL) and the antennal mechanosensory and motor centre (AMMC) (52). The ALs are the primary processing centers that receive all inputs conveying olfactory information gathered by the antennae (53), but also integrate inputs from other sensory modalities such as thermo, hygro and mechanoreception (54–56). Here we have shown that axons projecting from GRNs housed in the taste sensilla of the distal flagellomere innervate the ALs. Notwithstanding we have not found arborizations in other brain regions, we cannot discard that antennal GRNs other than those studied here can arborise in other brain regions. Mass backfills of *R. prolixus* antennae have shown axons projecting to the ALs and AMMC but also to the subesophageal (SOG), prothoracic and posterior ganglia (57). The SOG is commonly known as the brain processing center for gustatory information (58), receiving inputs mainly from mouthpart sensilla but also from gustatory neurons present in other appendages (59, 60). For example, antennal GRNs in moths and bees project directly to the SOG and the AMMC (59, 61–64). However, the SOG also receives axons from olfactory receptor neurons of the proboscis labellum in the mosquito *Anopheles gambiae* (65). Even if the SOG and ALs are known as a gustatory and an olfactory center, respectively, our results and those of Riabinina *et al*. (65) indicate that these neuropils should not be considered exclusively dedicated to a single chemosensory modality. Considering that different sensory modalities reach the ALs (*e.g.* olfactory, temperature, humidity, mechanosensory and gustatory), this neuropile should indeed be considered as multimodal brain processing center of antennal information in insects. Therefore, the hypothesis that ALs act as multimodal units deserves to be further investigated.

## Conclusions

The detection of aversive taste stimuli induces avoidance responses in animals. Compounds activating aversive responses, eliciting the avoidance of inedible food or preventing oviposition on inappropriate sites, are considered deterrent (66). Those that reduce the rate of biting of blood-feeding insects are of particular interest due to the inherent risk posed by each bite. The use of repellents is currently the main method employed worldwide to prevent nuisance bites from hematophagous insects (34). While olfaction is one of the main targets of most repellents used so far, the taste sense should not be disregarded. If an olfactorily-based preventive barrier is overcome, insects would land over the host and likely feed on it. Knowing that the taste sense drives rejection may allow finding gustatory deterrents for disease-vector insects complementing known repellents. High-salt avoidance in different contexts could be valuable as a method to control biting/feeding and enhancing avoidance responses in a disease vector that affects millions of people (33). Our findings open novel targets and strategies to prevent biting and feeding, as well as to promote substrate avoidance in a disease vector.

## Materials and Methods

### Insects

Fifth-instar larvae and adults of *R. prolixus* were used throughout the experiments. Insects obtained from the laboratory colony were reared at 28°C, ambient relative humidity (RH) and 12h: 12h L/D cycle. Fifth instars or adult insects used in the experiments did not have access to food following ecdysis. Fifth-instar larvae were used for feeding assays and RNAi experiments 7 - 21 days post-ecdysis. Adults used for neuroanatomy experiments were 7 - 9 day-old post-ecdysis.

### Feeding assays

Feeding behavior was examined using an artificial feeder described earlier (31). Briefly, it consisted of a recipient (*i.e.* the feeder) filled up with the appetitive solution (AS) with its lower opening closed with a latex membrane (*i.e.* bite substrate mimicking the host skin). The AS consisted of a 0.001 M aqueous solution of adenosine 5’-triphosphate disodium salt hydrate (ATP) (Sigma Aldrich, St. Louis, US) and 0.15 M NaCl (Biopack, Buenos Aires, AR). Insects were placed individually inside a plastic vial. A piece of filter paper placed vertically inside the plastic vial allowed bugs to reach the membrane of the feeder. The feeder containing the AS was placed close to an aluminum plate connected to a thermostatized resistance that heated the AS to 35 °C. The membrane in contact with the solution acquired the same temperature. The heat emitted by the recipient motivated the insects to approach the heated source (31, 67, 68).

Insects were weighed individually before (initial weight, Wi) and after (final weight, Wf) the feeding assay. Assays started when both containers, the plastic vial with the insect and the feeder with the heated AS, were put in contact and lasted for 10 minutes. Bugs had to pierce the membrane with their mouthparts in order to ingest the AS. A normalized weight gain was calculated as: (Wf - Wi) / Wi. Insects that ingested at least one time their own initial weight were considered as fed (31). Consequently, the percentage of insects whose normalized weight gain was higher than 1 was calculated. Experiments were carried out at the beginning of the scotophase, when kissing bugs display their maximal motivation to feed (69–71).

### Effect of salt concentration

Before reaching the AS insects had to get in contact with the treated membrane, which was impregnated with 50 μl of NaCl (Na) or KCl (K) at the following concentrations 0.15 M (low-Na/low-K), 0.5 M (mid-Na/mid-K), 1 M (high-Na/high-K) or with distilled water (no-Na/no-K, as control). These concentrations were chosen based on the amount of NaCl present on the human sweat, *i.e.* 0.1 − 0.17 M (24).

### Effect of amiloride

Amiloride hydrochloride hydrate (AMI) (Sigma Aldrich, St. Louis, US) was applied by allowing insects to walk freely for 1 minute over a filter paper (3 cm diameter) coated with 50 μl of 0.001 M AMI (+AMI). Thus, when the insect walked over the treated substrate the antennae as well as other body parts were exposed to AMI. The control group (−AMI) consisted of insects exposed to a filter paper impregnated with distilled water. During the following 3 minutes, insects were transferred to the artificial feeder paradigm in which responses to the same stimuli tested in the initial experiment were recorded.

### Role of the antenna

The last third of the distal flagellomeres of both antennae was cut off 24 h prior to the assays. Feeding performance was tested with bugs from the ablated group (−ANT) and intact ones (+ANT) as control. As previously, the membrane of the artificial feeder was coated with 50 μl 1 M NaCl (high-Na) or with distilled water (no-Na).

### Data analysis

Data were analyzed statistically by means of contingency tables of independence (Figure 1) (72). A global comparison including all treatments was assessed by means of a Pearson’s Chi-squared test (*X*^2^). Then, whenever the global test was statistically significant (α = 0.05), *post hoc* comparisons were carried out. To avoid experiment-wise error, the α value was corrected with the Bonferroni correction (α’ = α / k, k = number of comparisons) (72). The standard errors of percentages were calculated as √ p (1 – p) / N; p: proportion of response; N: number of animals tested (72).

### Electrophysiological recordings

Single sensillum recordings (73) were carried out on the 4 most distal taste sensilla of the distal flagellomeres, previously characterized by Pontes *et al*. (31) (shown in Figure 2A). Insects were immobilized inside a plastic pipette, with both antennae kept outside and fixed. Insects were grounded *via* insertion of a Ag/AgCl wire to the anus (reference electrode), and a glass capillary (20 μm in diameter at the tip) holding a Ag/AgCl wire inside served for both stimulus presentation and GRN response recording. Stimulus presentation lasted for 2 seconds and started once the glass capillary covered the tip of a single sensillum. The interval between subsequent stimulus presentations was 2 minutes. The biological signals were amplified, filtered (preamplifier: gain ×10, TastePROBE DTP-02, Syntech, Kirchzarten, DE; amplifier: gain × 100, eighth-order Bessel, pass-band filter: 10 – 3000 Hz, Dagan Ex1, Minneapolis, US), digitized (sampling rate: 10 kHz, 16 bits, Data acquisition module DT9803, Data Translation, Massachusetts, US) and stored in a PC. Spike detection and sorting was performed off-line using the dbWave software (74). Spike frequencies were calculated as the number of spikes *per* 1 second for each GRN.

### Physiological responses of GRNs

Taste sensilla were stimulated with 0.0001, 0.001, 0.01, 0.1, 0.2 and 0.5 M NaCl or KCl, Spike frequencies were calculated for each concentration. Additionally, the firing profile over time to 0.01 M NaCl/KCl was characterized for the different GRN types. To do this, the number of spikes *per* 100ms bin was calculated across the 2 seconds of each recording/GRN and averaged for neurons of the same type. Peristimulus histograms were built to visually analyze the temporal firing pattern of the two GRN types identified.

### Effect of amiloride on GRNs

Taste sensilla of the distal flagellomeres (Figure 2A) were applied with 0.001 M AMI by gently touching them with an impregnated toothpick. This procedure allowed us to approach the protocol applied for behavioral experiments with AMI (see before). Two control groups were required, one group with sensilla applied with a toothpick coated with distilled water (−AMI) and the other with sensilla applied with a dry toothpick (MCt). The latter being a mechanical control for the procedure. It is expected that touching the sensilla with a toothpick coated with distilled water (solvent used for AMI) or kept dry would have no effect on the physiology of GRNs, as shown in Figure S1. Afterwards the firing responses of GRNs to 0.01 or 0.2 M NaCl or KCl were recorded. The effect of amiloride was evaluated over the same sensilla along time. For this, GRN firing responses to 0.01 M NaCl/KCl were recorded at 0, 10, 20 and 30 minutes post-treatment.

### Physiological responses of GRNs after gene silencing

The firing activity of GRNs to 0.01 M NaCl and KCl was recorded for non-injected and ds-RNAi injected insects (see below).

### Data analysis

The effect of AMI on the firing frequency of the GRNs was statistically analyzed using a one tailed Wilcoxon matched-pairs signed rank test to compare responses to the different salt concentrations (Figure 2F-2G) or post-application times (*i.e.* 0, 10, 20, 30 min) (Figure 2H-2I). To avoid experimental-wise error, the α value (0.05) was corrected with the Bonferroni correction in Figure 2H-2I, where *k* = 4 and α’ = 0.0125 (72). The effect of gene silencing was assessed by using Kruskal-Wallis test followed by *post hoc* Dunn’s comparisons on the responses of GRNs to salts.

### Tracing of GRNs

Two neuronal tracers were used: neurobiotin (1% in 0.25 M KCl, Neurobiotin Tracer®, Vector Laboratories, Burlingame, US) and rhodamine (1% in distilled water, Dextran, Tetramethylrhodamine, 3000 MW, Anionic, Lysine Fixable, Thermo Fisher, Buenos Aires, AR). Neurobiotin and rhodamine were applied in the 4 distal taste sensilla of the distal flagellomeres (Figure 2A). Independently of the tracer used, the tip of the taste sensilla was previously cut to allow the dye to be incorporated into the gustatory neurons. Thereafter, cut sensilla were immersed for 6 minutes in distilled water. Neurobiotin was applied in a single sensillum, whereas rhodamine was applied simultaneously in the 4 taste sensilla, by inserting a single sensillum or the 4 sensilla in glass capillaries containing the corresponding dye.

To assure that only GRNs were stained, the two antenna of same animal were applied simultaneously with the rhodamine (the tip of each antenna was placed inside individual glass capillaries). However, while the 4 taste sensilla of one antenna were cut, the sensilla of the other antenna were kept intact.

Neurobiotin and rhodamine were allowed to migrate to the brain for 12h at room temperature or for 48h at 4°C, respectively. Following dye incubation times, the brains were dissected in Millonig’s buffer and fixed in 4 % paraformaldehyde overnight at 4°C. Neurobiotin brains were dehydrated and rehydrated in an alcohol series (50 %, 70 %, 90 %, 100 %) and propylene oxide and next incubated in Oregon Green-avidin (Oregon green^®^ 488 conjugate, Molecular probes, Oregon, US) with 0.2 % Triton X and 1 % BSA overnight at 4 ° C (58). Subsequently, neurobiotin brains were rinsed in Millonig’s buffer and dehydrated in the alcohol series. Rhodamine treated brains were rinsed in Millonig’s buffer and dehydrated through sequential baths in the same ethanol series and propylene oxide. Then, all the brains were cleared and mounted in methyl salicylate. Whole mounts were optically sectioned and scanned with a laser scanning confocal microscope (Olympus FV300/BX61, Carl Zeiss NTS SUPRA 40, Centro de Microscopía Avanzada, Facultad de Ciencias Exactas y Naturales, Universidad de Buenos Aires).

### Antennal expression of PPKs

One hundred distal flagellomeres from fifth instar larvae were excised, and a total of three replicates were prepared. Samples were manually homogenized using sterilized pestles and total RNA was extracted using 500 μL of TRIzol^®^ Reagent (Life Technologies, Carlsbad, US) according to the manufacturer’s instructions. The concentration of RNA extraction products was determined using a Qubit^®^ 2.0 fluorometer (Life Technologies, Carlsbad, US). All samples were treated with RQ1 RNase-Free DNase (Promega, Fitchburg, US). Treated RNA (11 μL per sample) was used to produce cDNA by means of the SuperScript III Reverse Transcriptase (Life Technologies, Carlsbad, US) and a 1:1 mix of Random Hexamers and 10 μM Oligo(dT) 20 primers to a final volume of 20 μL. For qPCR reactions, 7.5 μL of FastStart SYBR Green Master (Hoffmann-La Roche, Basel, SW) were used in the reaction that also contained 0.3 μL of a 10 μM primer solution and 1 μL of cDNA sample into a final volume of 15 μL. The reactions were conducted using the Mini Opticon Real-Time PCR Detector Separate MJR (Bio-Rad Laboratories, California, US) under the following conditions: 10 min cycle at 95°C, followed by 40 cycles of 20 s at 95°C, 20 s at 60°C and 30 s at 72°C. Real-time data were collected through the CFX Manager 3.0 software (Bio-Rad Laboratories, California, US). After qPCR reactions, melting curve analyses were performed to confirm reaction specificity. Non-template controls (NTC) were included for each primer set to verify the absence of exogenous DNA. To confirm amplicon sizes, qPCR products were run on a 2% agarose gel stained with ethidium bromide. NTCs were also included in the agarose gel.

### RNA interference

#### Double-strand RNA synthesis

Double strand RNAs (dsRNA) of *RproPPK28* and *RproPPK014276* were synthesized by amplification of antennal cDNA by means of PCR. The ampicillin resistance beta-lactamase gene (β-lact) of *Escherichia coli* was also amplified from the pBluescript SK plasmid as control of dsRNA injection (see below). PCR was carried out by using specific primers conjugated with 20 bases of the T7 RNA polymerase promoter (Table S1). PCR products, 200, 202 and 253 bp for *RproPPK28* and *RproPPK014276* and β-lact, respectively, were used as templates for dsRNA synthesis using the MEGAscript™ RNAi Kit (Thermo Fisher Scientific, Massachusetts, US). After synthesis, the purity and integrity of dsRNA were confirmed by running a 1.5 % agarose gel and quantified using NanoDrop™ (Thermo Fisher Scientific, Massachusetts, US).

#### ds-RNA injection

Insects were randomly separated in four experimental groups for dsRNA injections. Two groups of insects were injected with the dsRNAs against the genes of interest: dsRNAi-*RproPPK014276* and dsRNAi-*RproPPK28*. A third group was injected with the dsRNA of β-lact, representing a control for dsRNA injection: dsRNAi-ctrl. A microliter syringe (World Precision Instruments, Florida, US) was used to inject 2 μl of 1.25 μg/μl dsRNA diluted in PBS1X into the thoracic hemolymph of insects. The fourth group of insects was maintained intact (non-injected group) to control for potential effects of the injection procedure. Eleven days after dsRNA injection, insects of each group were tested for qPCR verification of gene expression knock-down, in electrophysiological recordings or in behavioral assays (see below).

To evaluate the efficacy of PPK knock-down, 20 antennae *per* group were excised (6 replicates per treatment). Total RNA was extracted as previously described using 250 μL of TRIzol^®^ Reagent, treated with DNAseI (Promega, Fitchburg, US) and used to synthesize cDNA using the SuperScript III Reverse Transcriptase (Thermo Fisher Scientific, Massachusetts, US), to a final volume of 30 μL. Real time PCR reactions were performed as described for expression profile experiments, except for the instrument that was an AriaMx Real-Time qPCR System (Agilent, California, US). The relative gene expression was calculated using the 2 -ΔΔCt method (75).

First, the gene expression for each condition was normalized using the geometric mean of two reference genes that were previously reported to have stable expression in *R. prolixus* antennae, *RproGADPH* and *RproG6PDH* (76). Subsequently, the normalized expression of each gene was normalized using the expression calculated for the non-injected group.

#### Behavioral response after gene silencing

The two-choice preference arena used consisted in a rectangular acrylic box (10 cm × 5 cm) divided in two equal zones (25). The substrate (filter paper) of one zone was coated with 100 μl distilled water (no-Na/no-K), while that of the other zone received either 100 μl of 1 M NaCl (high-Na) or KCl (high-K). One insect was placed inside an inverted recipient at the center of the arena for 1 minute, in order to acclimate to the experimental condition. Following this time, the insect was released and its preference was recorded during 4 minutes using an infrared-sensitive video-camera connected to a digital recorder. A preference index (PI) was calculated as: the difference between the time spent at each zone of the arena divided by the total experimental time. PIs near 0 indicate lack of preference. PIs close to −1 or 1 show preference for either side of the arena. All experiments were carried out in a dark room during the scotophase (see above).

#### Data analysis

Relative expression levels of knock-down genes were statistically compared by using Kruskal-Wallis test followed by *post hoc* Dunn’s comparisons. For behavioral assays, the PIs obtained for each animal of non-injected and dsRNAi injected groups were statistically compared against a PI value 0 (*i.e.* no preference) by using one sample *t*-tests (25, 72).

## Acknowledgments

We thank Guillermo Pereyra for rearing and providing insects. We specially thank to Sheila Ons for providing equipment and PCR reagents. We deeply thank to Carolina Reisenman, Meghan Laturney, Sharon Hill, Rickard Ignell and Anupama Dahanukar and lab members for their fruitful comments and suggestions that improved the manuscript.

RBB, GP, JMLE, MLG, MBA are members of the CONICET (Consejo Nacional de Investigaciones Científicas y Tecnológicas), AC is a fellowship holder of the CONICET. The financial support for this research was provided by ANPCyT (Agencia Nacional de Promoción Científica y Tecnológica, grant code Préstamo BID PICT 2013-1253 to RBB, grant code Préstamo BID PICT 2015-2825 to GP and grant code Préstamo BID PICT 2016-3103 to JMLE). MGL thanks FIOCRUZ and INCT-EM (Project number: 573959/2008–0) for kindly supporting his research.

---

**Figure S1.**
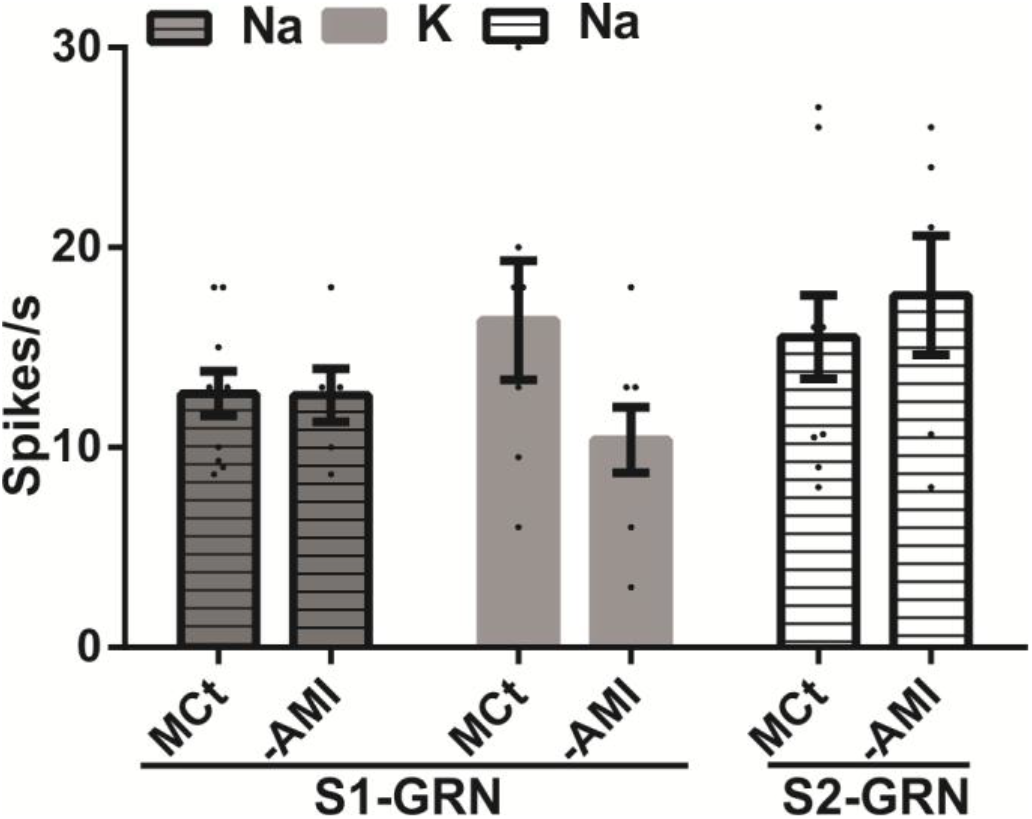
Effect of mechanical contact during amiloride application. Firing response of GRNs to Na or K following touching the taste sensilla with a toothpick kept dry (MCt) or coated with distilled water (−AMI). Scatter plots are shown and bars represent the mean of spike frequency (mean ± s.e.m) of GRNs in response to 0.01 M Na or K. No statistical differences were detected between MCt and -AMI groups (Wilcoxon test, MCt *vs.* −AMI, S1-GRN to Na: W = −3, p = 0.37 and to K: W = −10, p = 0.06; S2-GRN to Na: W = 4, p = 0.25, in all cases p > 0.05). Six to ten sensilla for each experimental condition were tested.

**Table S1.**
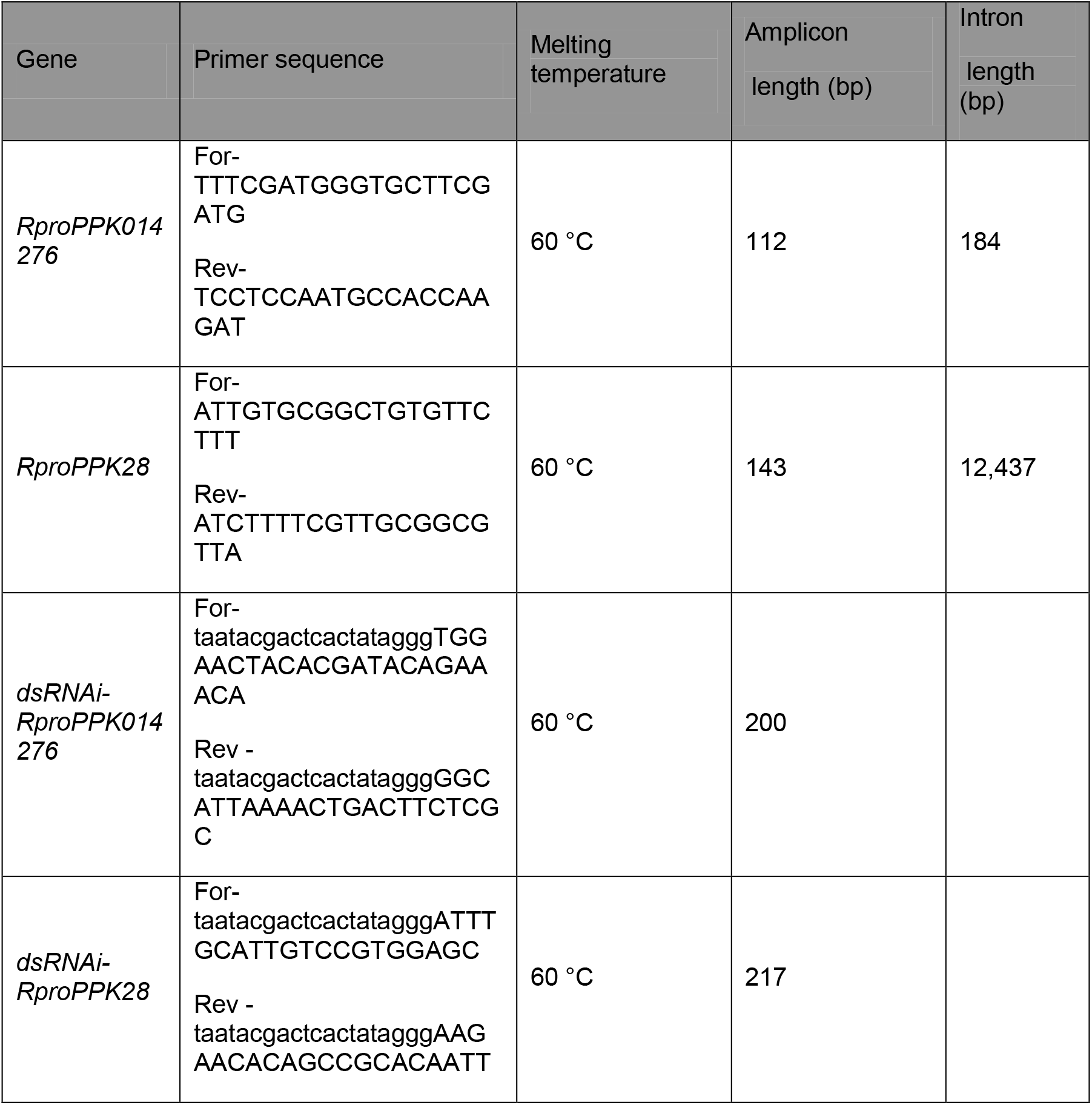
Specific primers used for qPCR and dsRNAi experiments.

